# Influenza A virus circumvents the innate immune response through the sequestration of double-stranded RNA

**DOI:** 10.1101/2024.05.22.595274

**Authors:** Masahiro Nakano, Sho Miyamoto, Chiho Ohnishi, Chiharu Nogami, Nanami Hirose, Yoko Fujita-Fujiharu, Yukiko Muramoto, Takeshi Noda

## Abstract

Double-stranded RNA (dsRNA), which induces an innate immune response against viral infections, is rarely detected in influenza A virus (IAV)-infected cells. Nevertheless, we previously reported that the IAV ribonucleoprotein complex (vRNP) generates looped dsRNAs during RNA synthesis *in vitro*. This finding suggests that IAV possesses a specific mechanism for sequestering dsRNA within infected cells, thereby enabling viral evasion of the innate immune response. Here, we found that dsRNAs were produced in infected cells lacking the expression of viral non-structural protein 1 (NS1) and nuclear export protein (NEP), both encoded by the same RNA segment. Interestingly, NS1 molecules masked the entire looped dsRNA generated by vRNP, implying a potential role for NS1 in segregating viral dsRNA from cytoplasmic dsRNA sensors, including retinoic acid inducible gene-I (RIG-I). Furthermore, dsRNAs were sequestered within the nucleus of infected cells due to the absence of NEP, while their translocation to the cytoplasm occurred only upon NEP expression. Notably, the cytoplasmic translocation of dsRNA triggered the innate immune response in an RIG-I-dependent manner. These findings highlight IAV’s distinctive strategy for circumventing innate immunity by sequestration of dsRNAs.

**Author summary:** It is widely recognized that double-stranded RNA (dsRNA) produced during viral infection triggers an innate immune response. However, the influenza A virus (IAV) has been thought to rarely produce dsRNA within infected cells. Here, we confirmed the limited dsRNA production in the nucleus of IAV-infected cells and found that the cells lacked expressions of viral non-structural protein 1 (NS1) and nuclear export protein (NEP), both derived from a single RNA segment. High-speed atomic force microscopy demonstrated that NS1 entirely concealed dsRNA produced by the viral ribonucleoprotein complexes, thereby segregating it from cytoplasmic dsRNA sensors that trigger the innate immune response. Interestingly, NEP expression caused cytoplasmic translocation of dsRNA, resulting in the nuclear translocation of interferon regulatory factor 3, which initiates an innate immune response. Collectively, our findings suggest that IAV employs a sophisticated strategy to circumvent the innate immune system, wherein the expressions of NS1 and NEP exert considerable influence.

## Introduction

It has long been acknowledged that double-stranded RNA (dsRNA) generated during viral infection induces an innate immune response [1–3]. In non-plasmacytoid dendritic cells, viral dsRNAs are recognized by cytoplasmic dsRNA sensors, such as retinoic acid-inducible gene-I (RIG-I) and melanoma differentiation-associated gene 5 (MDA5) [4,5]. These sensors have been reported to recognize distinct cytoplasmic dsRNA species; RIG-I detects 5’-triphosphate(ppp)-containing panhandle dsRNA, while MDA5 recognizes relatively long dsRNAs [6–11]. Upon recognizing viral RNAs, RIG-I and MDA5 change their conformation to form filamentous oligomers on substrate RNAs and associate with mitochondrial antiviral signaling protein (MAVS) [12–14]. The interaction between RIG-I/MDA5 and MAVS triggers downstream signaling pathway, ultimately resulting in the phosphorylation-driven nuclear translocation of interferon (IFN) regulatory factor (IRF)-3/7 and nuclear factor (NF)-κB. These activated factors orchestrate the transcriptional induction of type I and type III IFNs as well as proinflammatory cytokine genes [15–17]. While the majority of viruses generate dsRNA during their replication processes, it has been reported that certain negative-stranded RNA viruses, including influenza A virus (IAV), do not accumulate significant levels of dsRNA within virus-infected cells [18,19]. IAV has eight single-stranded viral genomic RNA segments (vRNAs), each of which is associated with multiple copies of a nucleoprotein (NP) and a heterotrimeric RNA-dependent RNA polymerase complex (PB2, PB1, and PA) to form a helical ribonucleoprotein complex known as the vRNP [20–22]. The transcription and replication of vRNA are conducted by the RNA polymerase complex in the context of vRNP in the nucleus of infected cells. Because nascent viral RNA is promptly separated from template vRNA following its synthesis [23], it is postulated that dsRNA intermediates are not produced during the replication process of IAV. Consequently, in IAV-infected cells, the induction of the innate immune response is thought to be triggered by the 5’-ppp panhandle structure of vRNA [11,24–26], defective viral genomes [27], or short aberrant vRNAs [28].

In contrast, we recently found that vRNP isolated from IAV virions frequently generates looped dsRNAs *in vitro* [29]. Notably, the vRNP associated with these looped dsRNAs underwent substantial deformation, adopting a nonhelical configuration. This phenomenon led us to speculate that dsRNA formation represents an aberration in RNA synthesis, in which nascent viral RNA fails to separate from the template vRNA. Additionally, we detected dsRNA in IAV-infected cells; however, the proportion of cells producing dsRNA was minimal (0.16% of infected cells) [29]. This finding suggests that IAV employs a specialized mechanism to sequester dsRNA within infected cells, thereby circumventing the innate immune response. In the present study, we aimed to determine how IAV suppresses dsRNA generation in infected cells, shedding light on the strategies employed to evade immune responses.

## Results

### IAV generates dsRNA in a limited number of infected cells

Previous studies indicated that IAV does not produce detectable amounts of dsRNA in infected cells [18,19]. In contrast, we previously found that the influenza A/Puerto Rico/8/34 (H1N1) (PR8) virus generated dsRNA within the nuclei of virus-infected Vero cells [29]. To assess dsRNA production across diverse cell lines, we infected various cell types with influenza A/WSN/33 (H1N1) virus at a multiplicity of infection (MOI) of 0.1. Immunofluorescence assays (IFA) using an anti-dsRNA antibody showed the absence of dsRNA in mock-infected cell lines (S1 Fig). In IAV-infected Vero cells, dsRNA was detected in the nucleus at 10 h post-infection (hpi) (Fig 1), in line with our previous study employing the PR8 virus [29]. Beyond 14 hpi, dsRNA was detected in the IAV-infected A549, HeLa, HuH-7, and 293T cells (Fig 1). Importantly, in these cell lines, dsRNA production was limited to a small number of infected cells, as previously documented in our study using Vero cells [29]. Contrary to our expectations, dsRNA was not detected in IAV-infected Madin-Darby canine kidney (MDCK) cells even at 24 hpi (Fig 1), suggesting that IAV did not generate detectable amounts of dsRNA in MDCK cells. These findings indicate that, while IAVs produce dsRNAs in various cell types, the production of dsRNA occurs under specific conditions.

**Fig 1.**
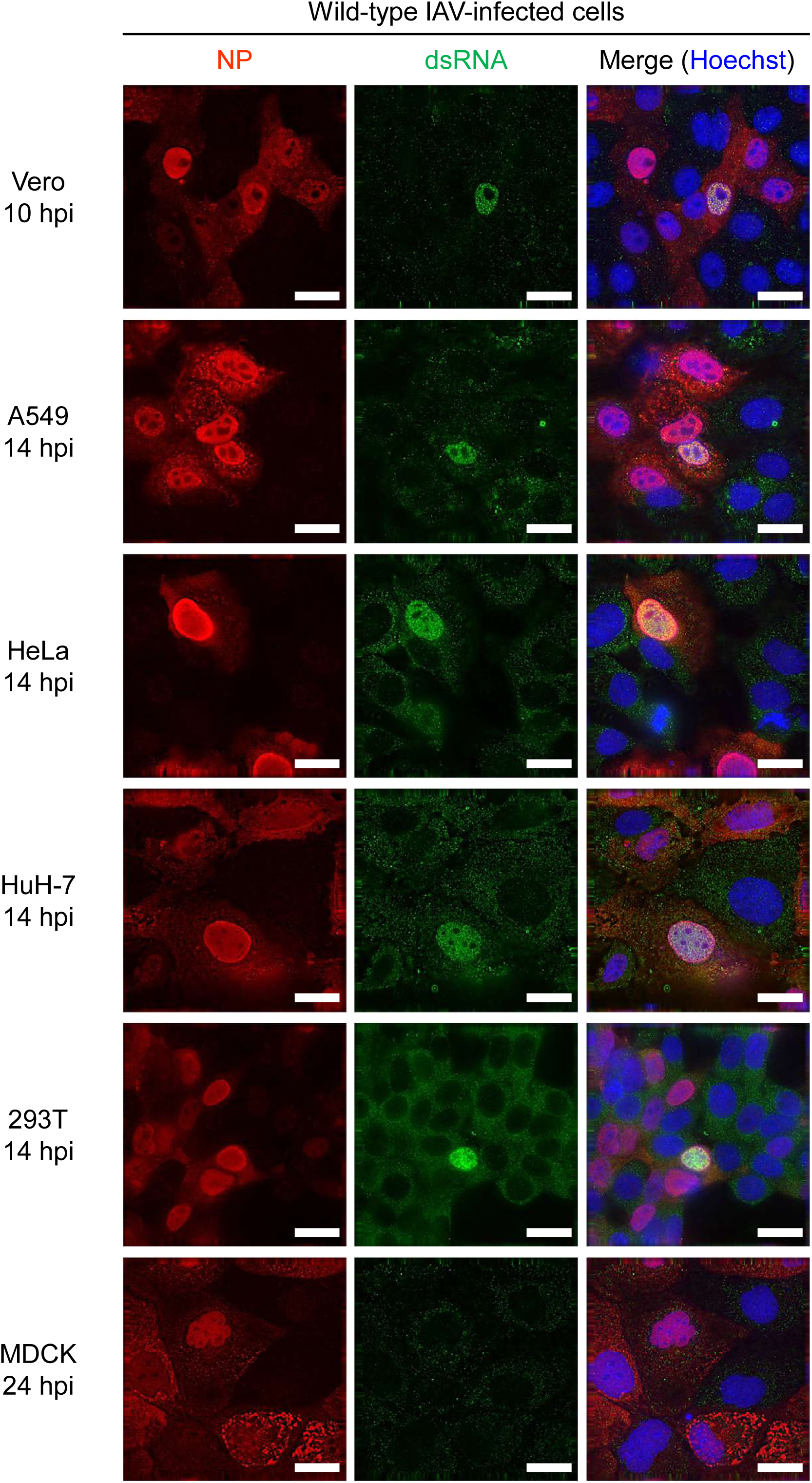
dsRNA production in IAV-infected cells. Cells were infected with the IAV WSN strain at an MOI of 0.1 and fixed at 10 hpi for Vero cells; 14 hpi for A549, HeLa, HuH-7, and 293T cells; or 24 hpi for MDCK cells. The presence of NP and dsRNA in infected cells was determined by IFA using anti-NP and anti-dsRNA antibodies, respectively. Cell nuclei were stained with Hoechst. The scale bars represent 20 µm.

### IAV produces dsRNA in infected cells lacking non-structural protein 1 (NS1) and nuclear export protein (NEP) expression

To understand the restrictive nature of dsRNA generation within IAV-infected cells, conditions for dsRNA production were explored. Vero and A549 cells were infected with IAV at different MOI, and the generation of dsRNAs was examined by IFA. Intriguingly, dsRNAs were barely observable at a high MOI (MOI = 5), whereas they were detected in infected cells at a low MOI (MOI = 0.1) (S2 Fig and S1 Table). Considering the established knowledge that IAV virions fail to express one or more viral proteins under low MOI conditions [30,31], we speculated that the expression of the viral protein responsible for preventing dsRNA generation was lacking in dsRNA-producing infected cells at a low MOI. Therefore, we investigated the relationship between dsRNA production and viral protein expression at a low MOI. Upon infecting A549 cells with IAV at an MOI of 0.1, NP and PA, which are integral components of vRNA replication, were ubiquitously expressed within the dsRNA-producing cells (Fig 2A, B, F and, S2 Table). Neuraminidase (NA), a spike protein present in the viral envelope, was detected in 50% of dsRNA-producing cells (Fig 2C, F and S2 Table). In contrast, two viral proteins originating from RNA segment 8, NS1 and NEP, exhibited limited expression in dsRNA-producing cells (Fig 2D–F and S2 Table). Simultaneous detection revealed the absence of NS1 and NEP in these cells (S3 Fig). These findings suggest that IAV generates dsRNA through vRNP during replication while concurrently suppressing dsRNA production via NS1 and/or NEP.

**Fig 2.**
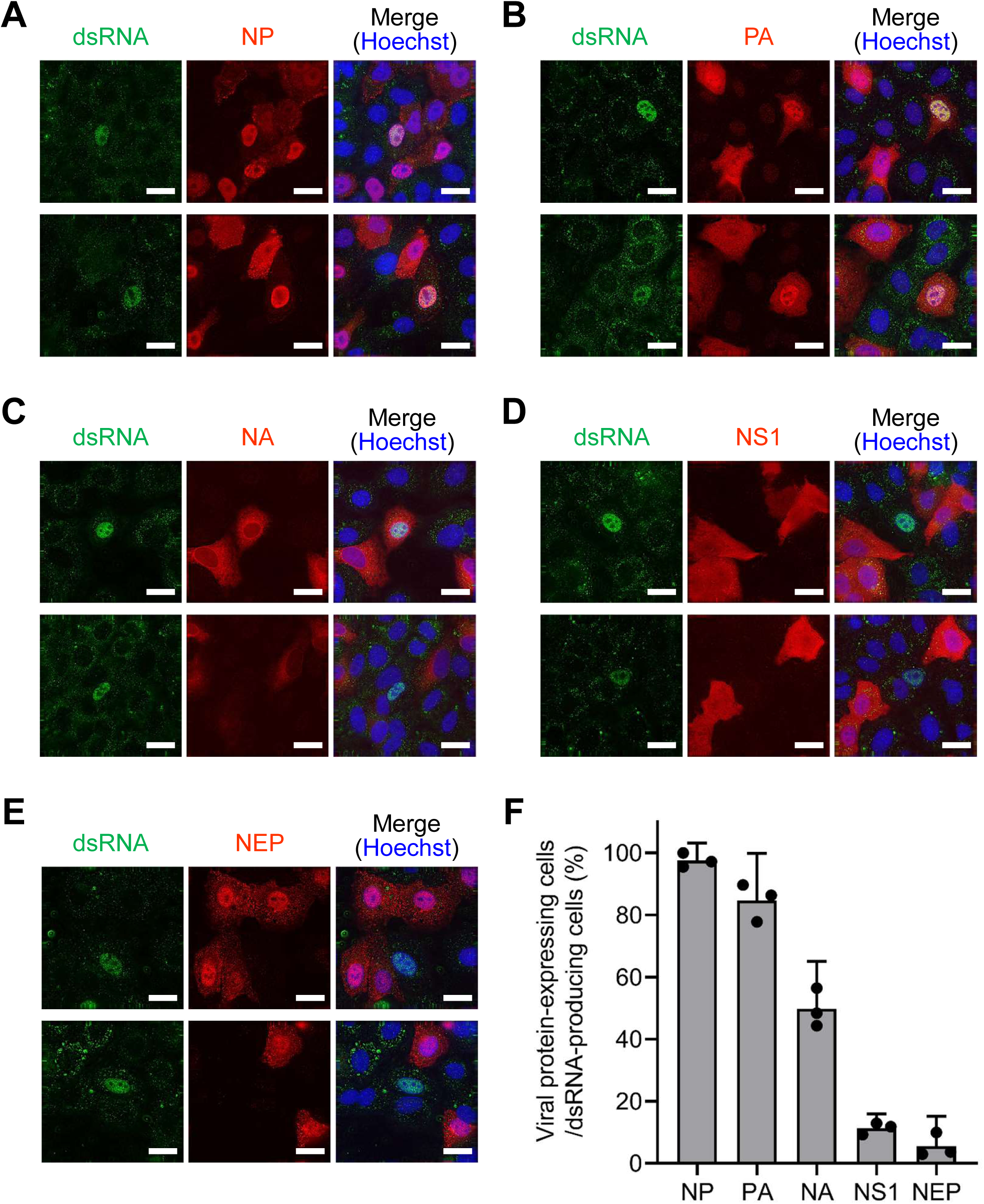
Viral protein expression in dsRNA-producing cells. A549 cells were infected with the IAV WSN strain at an MOI of 0.1 and subsequently fixed at 14 hpi. The presence of viral proteins in infected cells was detected by IFA using antibodies against NP (A), PA (B), NA (C), NS1 (D), and NEP (E). Additionally, an anti-dsRNA antibody was used to detect dsRNA, while Hoechst was used to stain the cell nuclei. The scale bars denote 20 µm. (F) The percentage of dsRNA-producing cells expressing each viral protein was calculated from the IFA results. The presented data represent the mean with 95% confidence intervals derived from three biologically independent experiments. Detailed counts of viral protein-expressing and dsRNA-producing cells are presented in S2 Table.

### IAV conceals looped dsRNA through the utilization of NS1

Because NS1 and NEP are absent in cells generating dsRNA, it was postulated that IAV suppresses dsRNA production through NS1 and/or NEP. NS1 acts as an antagonist of type I IFN and exhibits dsRNA-binding capabilities [32–34]. Accordingly, we focused on NS1 and prepared a mutant IAV lacking NS1 gene (ΔNS1 virus) [35] to examine the role of NS1 in inhibiting dsRNA production. As illustrated in Fig 3A, the ΔNS1 virus exclusively produces NEP from the truncated vRNA, while the wild-type IAV expresses both NS1 and NEP from the NS vRNA segment. To compare dsRNA production between ΔNS1 and the wild-type virus, Vero and A549 cells were infected with both viruses at the same MOI (MOI = 0.1), followed by IFA. The number of dsRNA-producing cells was 8–15 times larger in ΔNS1 virus-infected cells than in wild-type virus-infected cells (Fig 3B), implying that NS1 concealed dsRNA and prevented its detection in wild-type virus-infected cells. Interestingly, the ΔNS1 virus-infected cells exhibited a distinctive pattern; dsRNAs were detected not only in the nucleus but also in the cytoplasm of the infected cells (Fig 3C), which is different from the observation in wild-type virus-infected cells where dsRNA was detected only in the nucleus (Figs 1 and 2).

**Fig 3.**
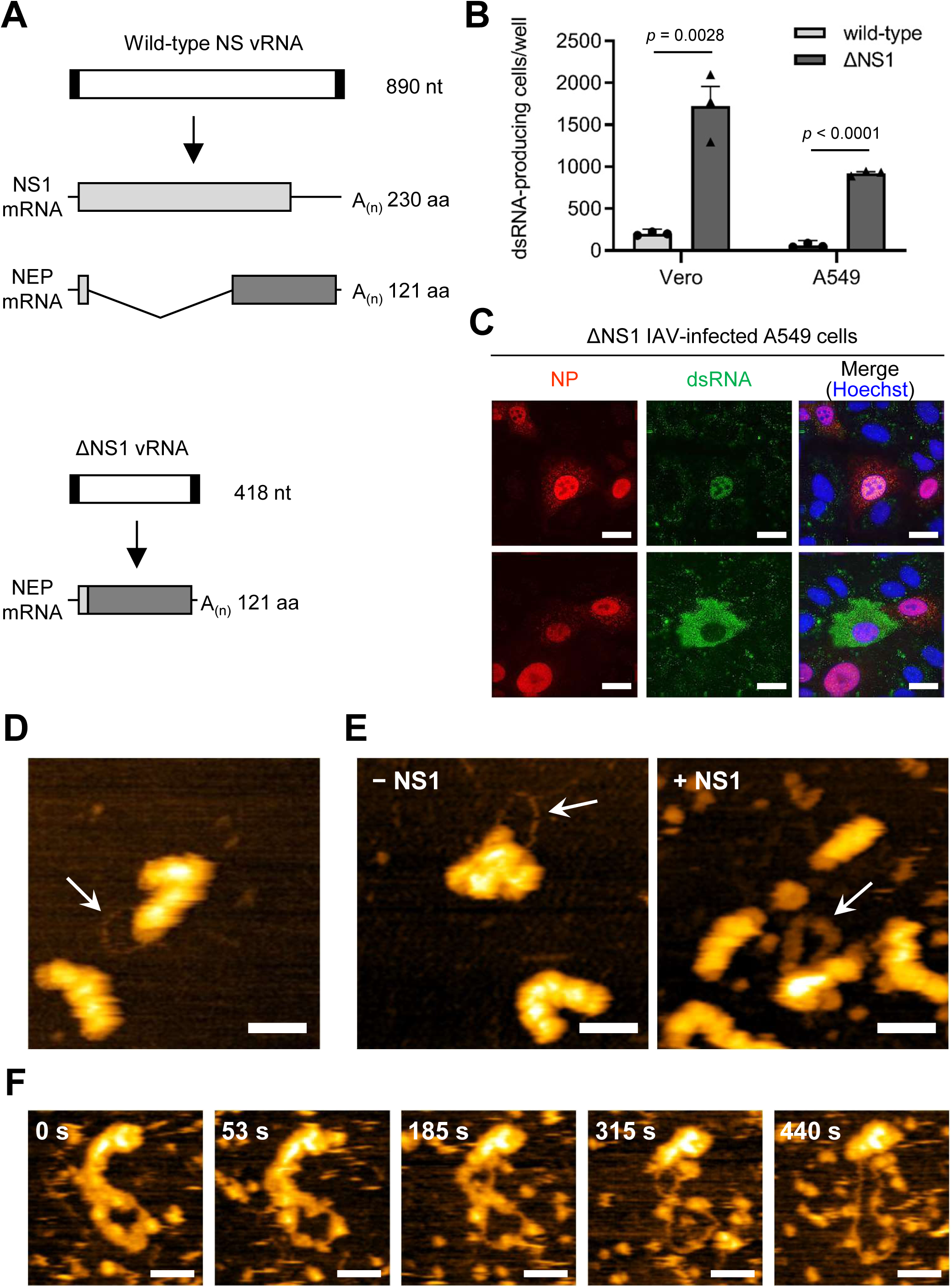
Concealment of dsRNA by NS1. (A) Schematic of NS genes and NS-specific mRNAs. A mutant NS gene which lacks NS1 open reading frame was designed, as described by Garcia-Sastre *et al*. [35]. Both NS1 and NEP mRNA were synthesized from wild-type NS vRNA (upper panel), whereas only NEP mRNA was transcribed from ΔNS1 vRNA (lower panel). The vRNA segments are depicted as white boxes flanked by black boxes, which denote the non-coding regions of the gene. The coding sequence of the NS1 protein is shown as a light gray box, whereas that of NEP is shown as a dark gray box. NEP mRNA originating from the wild-type NS vRNA is a spliced product of the NS1 mRNA, as shown by the V-shaped line. (B) Comparison of dsRNA production in wild-type IAV and ΔNS1 virus infections. Vero and A549 cells were infected with wild-type IAV or ΔNS1 virus at an MOI of 0.1 and were fixed at 24 hpi. Infected cells producing dsRNA were counted by IFA with an anti-dsRNA antibody. The presented data reflect the average ± 95% confidence intervals from three biologically independent experiments. Significance was determined using multiple unpaired *t*-test. (C) dsRNA production in cells infected with the ΔNS1 virus. A549 cells were infected with the ΔNS1 virus at an MOI of 0.1 and were fixed at 14 hpi. The presence of NP and dsRNAs was detected via IFA using anti-NP and anti-dsRNA antibodies, respectively. Cells producing dsRNAs within the nucleus (upper panels) and cytoplasm (lower panels) are shown. Cell nuclei were stained with Hoechst. Scale bars: 20 µm. (D) HS-AFM observation of vRNPs purified from FLAG–PB2 virus-infected cells. The presence of looped RNA associated with vRNP is indicated by an arrow. (E) Masking looped RNA via NS1. Following in vitro RNA synthesis using vRNPs purified from IAV virions, the recombinant NS1 protein was introduced. The mixture without (left panel) and with NS1 (right panel) was visualized using HS-AFM, with the looped structure in each image indicated by an arrow. (F) Detachment of NS1 from the looped RNA. The thick-looped structure observed after NS1 addition was monitored over an extended duration using HS-AFM, applying an augmented force. Five representative images captured at specific intervals are presented. All scale bars in the HS-AFM images are 50 nm.

Given that the vRNP purified from the IAV virion generated a looped dsRNA *in vitro* [29], we hypothesized that looped dsRNA is also generated in infected cells but is masked by NS1. To address this, the FLAG–PB2 WSN virus, which expresses the FLAG-tagged PB2 protein at its N-terminus [36], was produced using reverse genetics, and A549 cells were infected with this recombinant virus. At 24 hpi, FLAG-tagged vRNPs were obtained from the infected cells by immunoprecipitation and density gradient ultracentrifugation (S4A and B Figs). High-speed atomic force microscopy (HS-AFM) analysis revealed the presence of vRNP associated with looped RNA structures (Fig 3D), similar to those produced through *in vitro* RNA synthesis [29]. Furthermore, the looped RNA was digested by RNase III (S4C Fig and S1 Movie), indicating that the looped RNA observed in the infected cells was indeed dsRNA produced by the vRNP. However, contrary to our expectations, only naked-looped dsRNA was observed, and dsRNA masked by NS1 was not observed in the present study.

Subsequently, we investigated whether the recombinant NS1 protein could mask the dsRNA generated by the vRNP *in vitro*. Following *in vitro* RNA synthesis using purified vRNP, the purified recombinant NS1 protein (S5 Fig) was introduced, and the mixture was analyzed by HS-AFM. Remarkably, upon NS1 addition, thick looped structures were observed (Fig 3E and S6 Fig). The looped structures had lower heights (∼3.5 nm) than the vRNP and displayed uniform widths (∼25 nm) (S6 Fig). Prolonged HS-AFM observations (∼5 min) with the application of increased force revealed that individual particles sequentially detached from the looped structure, exposing the naked RNA (Fig 3F and S2 Movie). These findings strongly indicate that NS1 masks looped dsRNA generated by vRNP, a phenomenon that likely occurs within infected cells.

### IAV dsRNA undergoes translocation to the cytoplasm mediated by NEP

As shown in Fig 3C, within the ΔNS1 virus-infected cell, dsRNA was detected not only in the nucleus but also in the cytoplasm. Considering that the looped dsRNA associated with vRNP was isolated from infected cells (Fig 3D), we hypothesized that the cytoplasmic detection of dsRNA originated from the initial generation of dsRNA within the nucleus, followed by subsequent translocation to the cytoplasm. To investigate the potential nuclear export of dsRNA, we sought to distinguish between cells that generate dsRNA in the nucleus and those that generate dsRNA in the cytoplasm. The expression of viral proteins in ΔNS1 virus-infected dsRNA-producing A549 cells was also examined. When dsRNA was detected in the nucleus, the expression of viral proteins, excluding NP, was verified in 25–53% of dsRNA-producing cells, indicating that these viral proteins — NA, M1, and NEP—were dispensable for dsRNA production (Fig 4A and S7 Fig). Conversely, when dsRNA was identified in the cytoplasm, the expression of the two viral proteins underwent substantial alterations; NEP and M1 were expressed in nearly all dsRNA-producing cells, whereas the expression of NA did not change significantly (Fig 4A and S7 Fig). Intriguingly, both NEP and M1 are known for their pivotal roles in the nuclear export of vRNP [37–39], suggesting the possibility that dsRNAs are transported from the nucleus to the cytoplasm along with vRNP, mediated by these viral proteins.

**Fig 4.**
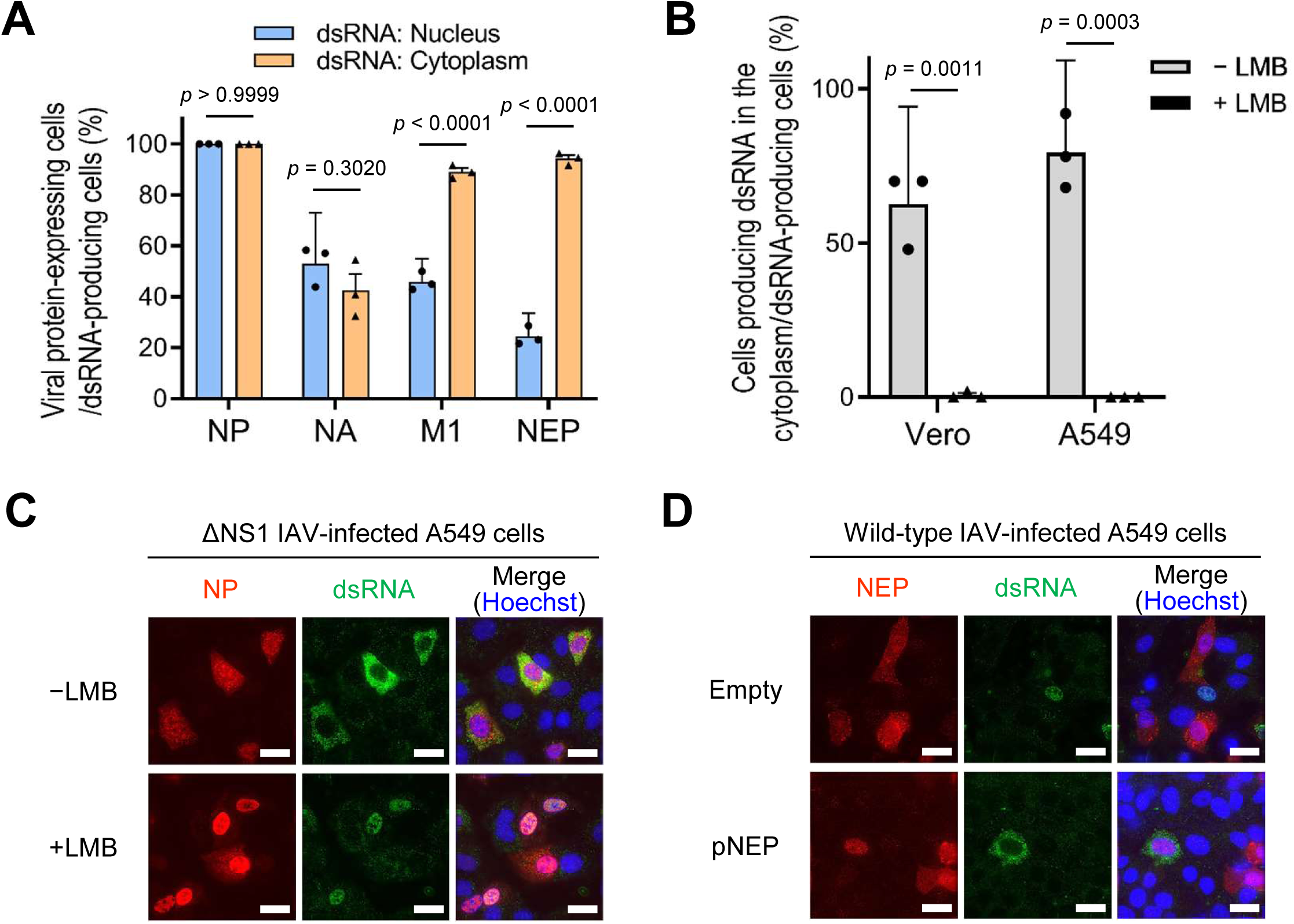
Cytoplasmic translocation of dsRNA mediated by NEP. (A) Relationship between dsRNA localization and viral protein expression in ΔNS1 virus-infected cells. A549 cells were infected with ΔNS1 virus at an MOI of 0.1 and fixed at 14 hpi. The production of dsRNA and expression of each viral protein were detected by IFA, as illustrated in S7 Fig. Percentages of cells expressing each viral protein in cells producing dsRNA within the nucleus (blue) or in the cytoplasm (orange) were calculated from the IFA results. Approximately 50 dsRNA-producing cells were examined in each experiment, and the data represent the average ± 95% confidence intervals of three biologically independent experiments. Significance was determined using two-way analysis of variance with Tukey’s test. (B and C) Inhibition of dsRNA nuclear export by LMB. Vero or A549 cells were infected with ΔNS1 virus at an MOI of 0.1 and subsequently subjected to incubation with or without LMB. The cells were fixed at 24 hpi, and dsRNA was detected by IFA. The subcellular localization of dsRNA was examined both in the absence (grey) and presence (black) of LMB. The computation of the proportions of cells producing dsRNA in the cytoplasm relative to the entirety of dsRNA-producing cells is delineated in B. In each experiment, approximately 50 cells generating dsRNA were examined, and the presented data delineate the mean ± 95% confidence intervals acquired from three biologically independent experiments. Significance was determined by multiple unpaired *t*-test. Representative A549 cell images obtained by IFA employing anti-NP and anti-dsRNA antibodies are displayed in C. Scale bars: 20 µm. (D) Cytoplasmic translocation of dsRNA in wild-type IAV-infected cells mediated by the expression of NEP. A549 cells were infected with wild-type IAV WSN strain at an MOI of 0.2 and were immediately transfected with the NEP expression plasmid (pNEP). The cells were fixed at 24 hpi, followed by staining with anti-NEP and anti-dsRNA antibodies. As a control, the pCAGGS vector was transfected (Empty). Scale bars: 20 µm.

Considering that dsRNA appears to be transported to the cytoplasm through its association with vRNP, we investigated whether inhibiting vRNP nuclear export would diminish the cytoplasmic localization of dsRNA. Vero and A549 cells were infected with the ΔNS1 virus in the presence or absence of leptomycin B (LMB), a known inhibitor of the nuclear export of vRNP via CRM1 [40], and the localization of dsRNA was assessed using IFA. In the absence of LMB, approximately 60% and 80% of dsRNA-producing Vero and A549 cells, respectively, exhibited cytoplasmic localization of dsRNA (Fig 4B and C). In contrast, in the presence of LMB, cytoplasmic localization was significantly reduced in both Vero and A549 cells, with only approximately 1% of dsRNA-producing cells displaying cytoplasmic localization of dsRNA. These results strongly indicate that dsRNA is exported to the cytoplasm in association with the vRNP.

Subsequently, the involvement of NEP in nuclear export of dsRNA was examined using wild-type IAV. Given the concurrent absence of both NS1 and NEP in wild-type IAV-infected dsRNA-producing cells, as described above (S3 Fig), we speculated that, in wild-type IAV-infected cells, the detection of dsRNA was confined solely to the nucleus because of the absence of NEP. To address this, A549 cells were infected with wild-type IAV and transfected with the NEP expression plasmid, and the localization of dsRNA was evaluated using IFA. The results demonstrated that, through the expression of NEP, a robust localization signal of dsRNA was evident in the cytoplasm, whereas such signals were absent in cells transfected with an empty vector (Fig 4D). Altogether, these findings strongly indicated that dsRNAs, likely associated with vRNP, could undergo nuclear-to-cytoplasmic export mediated by NEP.

### IAV induces innate immune response upon the export of dsRNA to the cytoplasm

dsRNAs induce innate immune responses in virus-infected cells [1–3]. Our findings suggest that IAV dsRNA, upon translocation to the cytoplasm, is recognized by cytoplasmic dsRNA sensors such as RIG-I. Therefore, to ascertain whether IAV dsRNA was indeed responsible for eliciting the innate immune response, the production of dsRNA and nuclear translocation of IRF3 in each cell line were analyzed using IFA. Upon infecting A549 cells with wild-type IAV, dsRNAs were exclusively detected within the nucleus, and nuclear translocation of IRF3 was not associated with dsRNA production (S8 Fig). In ΔNS1 virus-infected cells, the nuclear translocation of IRF3 remained unaffected by dsRNA production when dsRNA was localized in the nucleus; the nuclear translocation of IRF3 was observed in approximately 40% of the cells producing dsRNA in the nucleus (Fig 5A, upper left, right). Conversely, when dsRNA was detected in the cytoplasm, nearly all dsRNA-producing cells exhibited nuclear translocation of IRF3 (Fig 5A, lower left, right). Similar results were observed for the nuclear translocation of NFκB (Fig 5B), suggesting that cytoplasmic localization of dsRNA induced the innate immune response, whereas nuclear localization of dsRNA did not necessarily elicit such a response. This observation was corroborated in wild-type IAV-infected cells. By transfecting the NEP expression plasmid into wild-type IAV-infected cells, dsRNA was localized in the cytoplasm, leading to nuclear translocation of IRF3 in these cells (Fig 5C).

**Fig 5.**
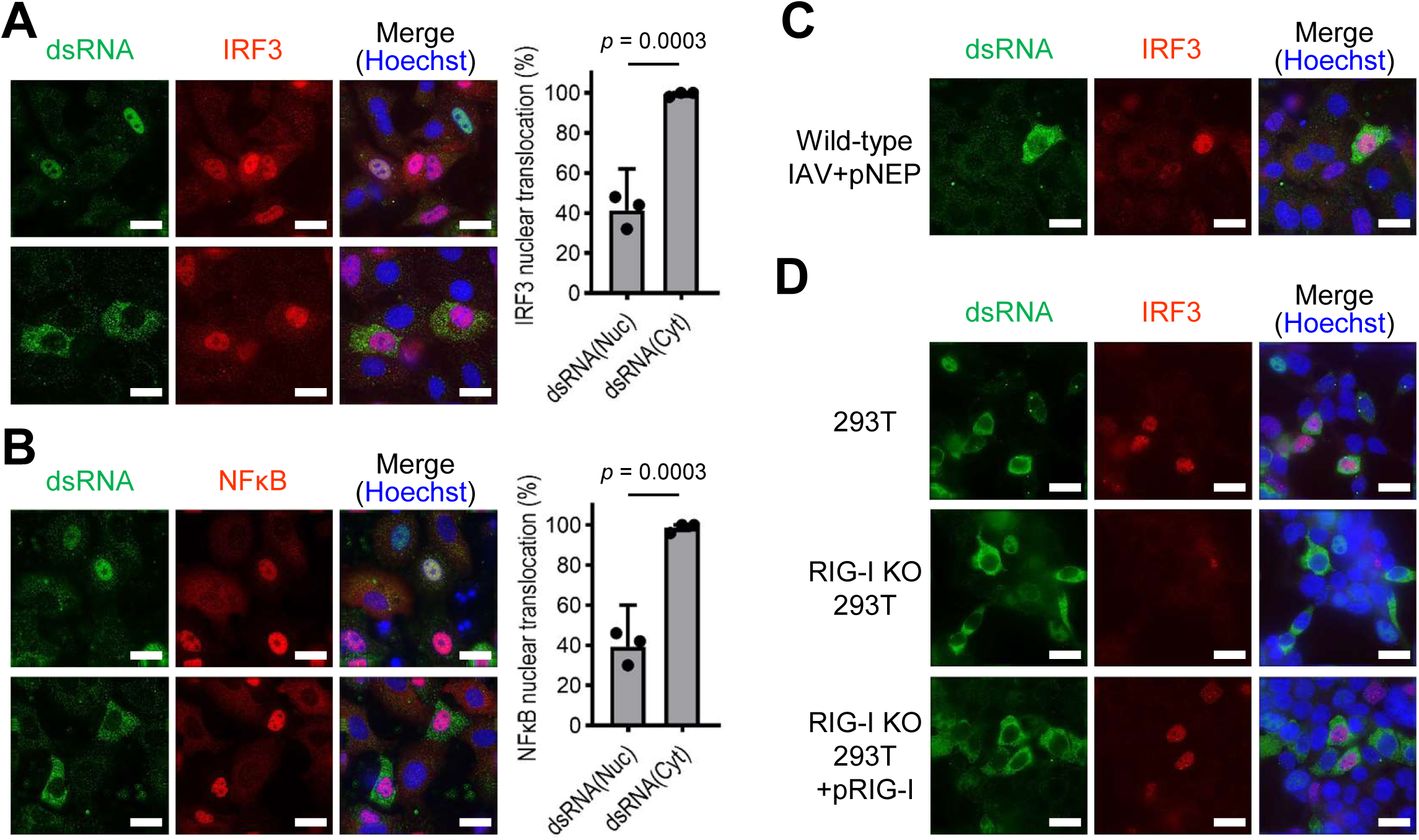
Induction of the innate immune response by IAV dsRNA. (A and B) Nuclear translocation of IRF3 and NFκB in dsRNA-producing cells. A549 cells were infected with ΔNS1 virus at an MOI of 0.2 and were fixed at 24 hpi. Antibodies against IRF3 (A) and NFκB (B) were employed in IFA for the identification of nuclear translocation of IRF3 and NFκB, respectively. The percentage of cells exhibiting nuclear translocation of each transcription factor within cells producing dsRNA in the nucleus (Nuc) or cytoplasm (Cyt) is shown on the right. For each enumeration, 50 dsRNA-producing cells were examined, and the data represent the mean ± 95% confidence intervals acquired from three biologically independent experiments. Statistical significance was determined by unpaired *t*-test. (C) IRF3 nuclear translocation in wild-type IAV-infected cells transfected with NEP expression plasmid. A549 cells were infected with wild-type IAV at an MOI of 0.2 and promptly transfected with the NEP expression plasmid (pNEP). Cell fixation was performed at 20 hpi, followed by staining with anti-dsRNA and anti-IRF3 antibodies. (D) RIG-I-dependent nuclear translocation of IRF3 in dsRNA-producing cells. Wild-type 293T (upper panels) and RIG-I KO 293T (middle and lower panels) cells were infected with the ΔNS1 virus at an MOI of 0.2. For the experiment involving the complementation of RIG-I functions, RIG-I KO 293T cells were transfected with the RIG-I expression plasmid at 2 hpi (lower panels). The cells were fixed at 24 hpi and stained with anti-dsRNA and anti-IRF3 antibodies. All scale bars denote 20 µm.

Finally, we assessed whether initiation of the innate immune response by dsRNA relies on the cytoplasmic RNA sensor RIG-I. For this purpose, we used RIG-I-knockout (KO) 293T cells [41,42]. Similar to A549 cells, nuclear translocation of IRF3 was observed in wild-type 293T cells, where dsRNAs were generated in the cytoplasm following ΔNS1 virus infection (Fig 5D, upper panels). Conversely, nuclear translocation of IRF3 was absent in ΔNS1 virus-infected RIG-I KO cells producing cytoplasmic dsRNAs (Fig 5D, middle panels), whereas this was confirmed in RIG-I KO cells transfected with poly(I:C), probably via MDA5 (S9 Fig). Through the transfection of the RIG-I expressing plasmid, the nuclear translocation of IRF3 became observable in the ΔNS1 virus-infected RIG-I KO cells generating cytoplasmic dsRNAs (Fig 5D, lower panels). These results suggest that the dsRNA generated by IAV infection can elicit an innate immune response in an RIG-I-dependent manner.

## Discussion

In the present study, we demonstrated that IAV generated dsRNA within virus-infected cells, wherein the expression of NS1 is notably absent. IAV appears to strategically utilize NS1 to conceal dsRNA, thereby segregating it from cytoplasmic dsRNA sensors such as RIG-I. Indeed, upon translocation into the cytoplasm via NEP, dsRNA derived from IAV induces the nuclear translocation of IRF3. These findings imply that IAVs employ a distinctive strategy to circumvent innate immune responses.

Although the occurrence of dsRNA-producing cells is limited, IAVs generate dsRNAs in diverse cell types (Fig 1). Previous studies have shown that dsRNAs are rarely detected in IAV-infected cells [18,19]. In these investigations, cells were infected with a relatively high MOI of IAV (MOI = 3–5), whereas we revealed the significance of a low MOI in facilitating dsRNA production (S2 Fig and S1 Table). IAV infection at a low MOI yields infected cells that lack one or more viral proteins [30,31]. Although the mechanism governing the generation of virions that express an incomplete set of viral proteins remains unclear, it appears to occur when virions fail to package one or more vRNA segments or when incoming vRNPs undergo unsuccessful nuclear import, transcription, and/or replication. IAV-infected cells expressing NP and RNA polymerases but lacking NS1 were unable to mask the dsRNA generated by vRNP, leading to the detection of dsRNA within the infected cells. Conversely, in cells expressing the complete repertoire of viral proteins, if dsRNAs were aberrantly generated during RNA synthesis, IAV would concurrently conceal them with NS1, thereby precluding the detection of dsRNA.

The absence of NS1 in dsRNA-producing cells led us to hypothesize that NS1 masks dsRNA expression in IAV-infected cells. It has been previously reported that NS1 forms a tubular structure in the presence of dsRNA which comprises complementary strands with GA and CU repetitions [34]; however, the capability of NS1 to mask dsRNA generated by IAV remains unrevealed. Therefore, we aimed to purify the looped dsRNA– vRNP complex from IAV-infected cells, wherein dsRNA was expected to be masked by NS1. However, solely the naked-looped dsRNA–vRNP complex was purified, and the looped dsRNA associated with NS1 was not observed (Fig 3D). This might be because NS1 molecules bound to dsRNA were dissociated during the purification process using detergent. In contrast, we observed the NS1 masking of looped dsRNAs produced by *in vitro* RNA synthesis. The looped dsRNA masked by NS1 exhibited regular width and height (Fig 3E and S6 Fig), suggesting that NS1 molecules were uniformly associated with the dsRNA. While NS1 molecules detached from the dsRNA during HS-AFM observation, they required a prolonged duration and substantial applied force for dissociation, implying that NS1 persistently masks dsRNA in infected cells. These findings suggest that IAV employs NS1 to conceal dsRNA, thereby evading the innate immune response.

We disclosed that IAV dsRNA induces the nuclear translocation of IRF3/NFκB upon the translocation of dsRNA to the cytoplasm (Fig 5), signifying the potential of IAV dsRNA to induce an innate immune response. Indeed, the absence of nuclear translocation of IRF3 in IAV-infected RIG-I KO cells substantiates the likelihood that IAV dsRNA is recognized by RIG-I in the cytoplasm. However, IAV has long been believed to not generate dsRNA, suggesting that the innate immune response is triggered by alternative RNA species. As RIG-I specifically recognizes the 5’-ppp moiety of panhandle dsRNA [11,24–26], the primary candidate for eliciting the innate immune response was considered to be full-length vRNA. Nevertheless, several studies have asserted that IFN induction requires viral RNA synthesis and that incoming vRNA is unlikely to be implicated in this process [43,44]. Consequently, the innate immune response is presumed to be prompted by aberrant RNA species that arise during RNA synthesis, such as defective viral genomes [27] and short aberrant vRNAs [28]. These RNA species harbor a 5’-ppp structure, making them recognizable by RIG-I and capable of inducing the innate immune response. The structural characteristics of the dsRNA identified in the current investigation, specifically whether it possesses a 5’-ppp structure, remains unknown. However, if the dsRNA represents a failure in vRNA replication, it should inherently bear a 5’-ppp structure, as replication occurs through *de novo* RNA synthesis. Indeed, the detectability of dsRNA during a later stage of infection, beyond the 14-h threshold, implies its emergence as a result of unsuccessful vRNA replication rather than transcription. Although further investigation is required, we presume that the dsRNA identified in the present study is a plausible candidate for triggering the innate immune response, supplementing an array of other RNA species reported previously.

Although IAV dsRNA seemingly possesses the potential to elicit an innate immune response, IAV ingeniously segregates dsRNA from cytoplasmic RNA sensors. The present study suggests that IAV masks the entirety of dsRNA with NS1. The translocation of this masked dsRNA to the cytoplasm remains uncertain; nevertheless, it will evade recognition by RNA sensors, given their incapacity to access the shielded dsRNA (Fig 6, upper). In contrast, in wild-type IAV-infected cells devoid of NS1 expression, detectable amounts of dsRNA are generated. However, in such instances, because both NS1 and NEP are encoded by the same RNA segment, with NEP expressed from the spliced NS mRNA (Fig 3A), NEP expression is simultaneously absent. Accordingly, in the context of wild-type virus infection, cells that lack NS1 but produce NEP would not be observed, causing the dsRNA to persist within the nucleus (Fig 6, lower). If NS1 and NEP are encoded by distinct RNA segments, a different situation may arise in which only NEP is expressed and dsRNA translocates to the cytoplasm. The simultaneous absence of NS1 and NEP results in sequestration of dsRNA within the nucleus, thereby segregating it from cytoplasmic dsRNA sensors. However, when NS1 incompletely masks dsRNA, NEP may transport unmasked dsRNAs to the cytoplasm. It has been documented that a singular dsRNA molecule alone possesses the capability to elicit the innate immune response [45]. Consequently, there is a potential scenario wherein undetectable amounts of dsRNA in the cytoplasm prompt the nuclear translocation of IRF3. For instance, in the current investigation, IRF3 nuclear translocation was also observed when dsRNA was detected in the nucleus, suggesting the possibility that this nuclear translocation was induced by undetectable cytoplasmic dsRNA (Fig 5A, upper left, right). Nevertheless, given that the proportion of IRF3 nuclear translocation when dsRNA was detected in the nucleus was less than 40%, it appears that IAV adeptly sequesters dsRNA from cytoplasmic dsRNA sensors by retaining it in the nucleus.

**Fig 6.**
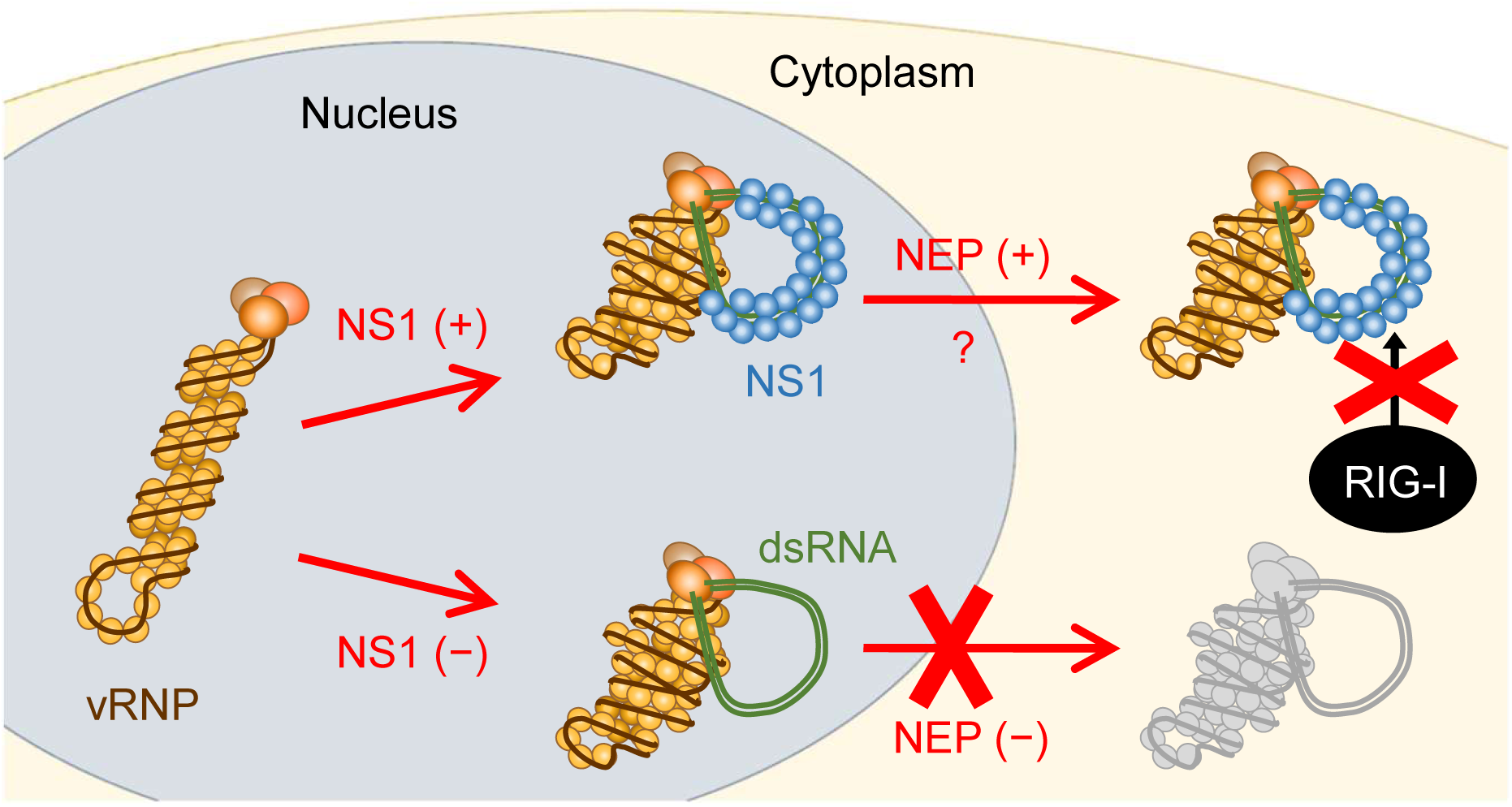
Proposed strategies for circumventing the innate immune response employed by IAV. IAV vRNP produces dsRNAs in infected cells. When NS1 is expressed (upper), dsRNA is concurrently masked by NS1. Although the translocation of masked dsRNA to the cytoplasm remains undetermined, RIG-I appears to be incapable of accessing IAV dsRNA, thereby enabling IAV to circumvent the innate immune response. Conversely, in scenarios in which NS1 expression is absent (lower), a detectable amount of dsRNA is generated within the nucleus. Nevertheless, in the absence of NS1 expression, NEP expression is also nonexistent, resulting in the sequestration of dsRNA within the nucleus and, consequently, the isolation of dsRNA from RIG-I.

Collectively, we propose a distinctive strategy employed by IAV to circumvent the innate immune responses. Considering that the induction of the innate immune response by dsRNA is ubiquitously observed across diverse viruses, our findings will exert influence not only on the investigation of IAV but also on other viruses.

## Materials and methods

### Cells

African green monkey kidney Vero cells (CCL-81; ATCC, Manassas, VA, USA) were grown in Eagle’s minimum essential medium (MEM; Gibco, Gaithersburg, MD, USA) supplemented with 10% fetal calf serum (FCS; Biosera, Nuaille, France). Human lung carcinoma A549 cells (CCL-185; ATCC), human cervical carcinoma HeLa cells (JCRB9004; JCRB Cell Bank, Tokyo, Japan), human hepatoma HuH-7 cells (JCRB0403; JCRB Cell Bank), and human embryonic kidney (HEK) 293T cells (CRL-3216; ATCC) were cultured in Dulbecco’s modified Eagle’s medium (DMEM; Merck, Darmstadt, Germany) with 10% FCS. MDCK cells were kindly provided by Y. Kawaoka (University of Tokyo) and grown in MEM supplemented with 5% newborn calf serum (Thermo Fisher Scientific, Waltham, MA, USA). HEK 293T RIG-I KO cells were kindly provided by T. Fujita (Kyoto University) and were grown in DMEM supplemented with 10% FCS. All cell cultures were maintained at 37°C in a 5% CO2 atmosphere.

### Plasmid construction

The NS1 deletion plasmid, denoted as pPolI-ΔNS1, was constructed following previously outlined procedures [35]. For the excision of the NS1 gene, an inverse PCR was performed, utilizing pPolI-NS1 (Provided by Prof. Kawaoka, University of Tokyo) as a template, along with a primer set: 5’-GACATACTGATGAGGATGTC-3’ (delNS1_WSN_F) and 5’-CTGAAAGCTTGACACAGTGTTTG-3’ (delNS1_WSN_R).

The resulting PCR product underwent DpnI treatment, purification through the MinElute PCR purification kit (Qiagen, Hilden, Germany), and subsequent ligation using T4 polynucleotide kinase (Takara, Kusatsu, Japan) and T4 DNA ligase (Ligation mix #6023; Takara). The pPolI-FLAG–PB2 plasmid was designed to generate FLAG–PB2 RNA, wherein the sequence encoding the FLAG peptide (DYKDDDDK) was fused to the sequence encoding the N-terminus of PB2 [36]. Construction of the plasmid involved the insertion of the FLAG–PB2 open reading frame with a stop codon into a truncated pPolI-PB2 plasmid with the 3’ and 5’ noncoding regions.

### Transfection

The NEP expression plasmid (pCAGGS-NEP) was transfected into A549 cells using Lipofectamine™ 3000 transfection reagent (Thermo Fisher Scientific). The RIG-I expression plasmid (pCAGGS-RIG-I) was transfected into wild-type or RIG-I KO 293T cells employing Polyethylenimine “Max” (PEI MAX; PolySciences, Warrington, PA, USA). The transfection of poly(I:C) (Invitrogen, Carlsbad, CA, USA) into wild-type or RIG-I KO 293T cells was conducted using the Lipofectamine™ RNAiMAX transfection reagent (Thermo Fisher Scientific). All transfections were performed according to the manufacturer’s instructions.

### Generation of recombinant viruses by reverse genetics

Reverse genetics was performed using pPolI plasmids encompassing the cDNA sequences of influenza A/WSN/1933 (WSN; H1N1) viral genes positioned between the human PolI promoter and the mouse PolI terminator, as previously outlined [46]. To generate wild-type IAV, eight pPolI plasmids expressing vRNA and four pCAGGS protein expression plasmids for PB2, PB1, PA, and NP were mixed with TransIT-293 (Mirus, Madison, WI, USA) and introduced into 293T cells. The cells were cultured in MEM supplemented with 0.3% bovine serum albumin (MEM/BSA) at 37°C. At 48 h post-transfection (hpt), the cells were treated with 1 µg/mL TPCK-trypsin (Worthington Biochemical, Lakewood, OH, USA) for 30 min, followed by centrifugation at 1,750 × *g* for 15 min at 4°C. The viral supernatant was collected and stored at −80°C. The FLAG– PB2 virus was generated by substituting pPolI-PB2 (wild-type) with the pPolI-FLAG– PB2 plasmid. For subsequent viral amplification, MDCK cells were infected at an MOI of 0.00001, followed by a 2-day incubation in MEM/BSA containing 1 µg/mL TPCK-trypsin. The ΔNS1 virus was generated by replacing pPolI-NS (wild-type) with the pPolI-ΔNS1 plasmid. Additionally, pPCAGGS-NS1 was co-transfected into 293T cells. The viral supernatant was harvested at 72 hpt and subsequently inoculated into Vero cells for viral replication. The inoculated cells were cultivated in MEM/BSA at 37°C for 5 days, with the periodic addition of 1 µg/mL of TPCK trypsin at 24-h intervals [47]. The resultant viral supernatant was harvested, and NS1 gene deletion was verified by reverse-transcription PCR and sequencing.

### IFA

Cells were cultured on a 35-mm glass-bottom dish (Matsunami Glass, Osaka, Japan) pre-coated with rat collagen type I (Corning, Corning, NY, USA) one day before infection. Subsequently, cells were infected with IAV (wild-type, ΔNS1, and FLAG–PB2 viruses) at an indicated MOI, followed by incubation for specified durations in MEM/BSA. In experiments concerning the inhibition of vRNP nuclear export, 5 ng/mL of leptomycin B (Cell Signaling Technology, Danvers, MA, USA) was introduced into the medium. After infection, the cells were fixed in 4% paraformaldehyde (Nacalai Tesque, Kyoto, Japan) for 10 min, followed by permeabilization with 0.1% Triton X-100 in phosphate-buffered saline (PBS) for 10 min. After washing with PBS, the cells were blocked with Blocking

One (Nacalai Tesque) for 30 min. Following the blocking step, cells were incubated with anti-NP rabbit polyclonal (1:1000 dilution, GTX125989; GeneTex, Irvine, CA, USA) and anti-dsRNA mouse monoclonal antibodies J2 (1:500 dilution, 10010200; Scicons; Nordic-MUbio, Susteren, The Netherlands) overnight at 4°C. Rabbit polyclonal antibodies against PA (1:1000 dilution, GTX125932; GeneTex), NS1 (1:1000 dilution, GTX125990; GeneTex), NEP (1:1000 dilution, GTX125953; GeneTex), NA (1:1000 dilution, GTX125974; GeneTex), and M1 (1:1000 dilution, GTX125928; GeneTex) were used instead of the anti-NP antibody to detect IAV proteins. For the assessment of nuclear translocation of IRF3 and NFκB, rabbit monoclonal antibodies against IRF3 (1:1000 dilution, 11904S; Cell Signaling Technology) and NFκB (1:1000 dilution, 8242S; Cell Signaling Technology) were employed instead of the anti-NP antibody, respectively. Following PBS washes, cells were incubated with Alexa Fluor 488-conjugated anti-mouse antibody (1:1000 dilution, A11001; Thermo Fisher Scientific) and Hoechst 33342 (Thermo Fisher Scientific) for 1 h at 4°C. After incubation, the cells were washed with PBS and incubated with Alexa Fluor 555-conjugated anti-rabbit antibody (1:1000 dilution, A21428; Thermo Fisher Scientific) for 1 h at room temperature. To simultaneously detect dsRNA, NP/NEP, and NS1, cells were incubated with Alexa Fluor 647-conjugated anti-NS1 mouse antibody (1:500 dilution, sc-130568 AF647; Santa Cruz Biotechnology, Dallas, TX, USA) for 1 h at room temperature subsequent to all incubation processes. All the antibodies were diluted in PBS containing 10% Blocking One. Sectional images were captured and deconvoluted using a DeltaVision Elite system (GE Healthcare, Chicago, IL, USA) with a 60× oil-immersion objective on an Olympus IX71 microscope.

### Purification of vRNP

The influenza A virus A/Puerto Rico/8/34 (H1N1) (PR8) was prepared according to previously reported procedures [48]. Purified PR8 virions (∼5 mg/mL) were lysed in a 50 mM Tris-HCl buffer (pH 8.0), encompassing 100 mM KCl, 5 mM MgCl2, 1 mM dithiothreitol (DTT), 2% Triton X-100, 5% glycerol, 2% lysolecithin, and 1 U/μL RNasin Plus RNase inhibitor (Promega, Madison, WI, USA) for 1 h at 30°C. Subsequently, the sample underwent ultracentrifugation through a 30% to 70% (w/v) glycerol gradient in Tris-NaCl buffer [50 mM Tris-HCl (pH 8.0) and 150 mM NaCl] at 250,000 × *g* for 3 h at 4°C, employing a SW55Ti rotor (Beckman Coulter, Brea, CA, USA). The collected fractions were mixed with 2× Tris-glycine sodium dodecyl sulfate (SDS) sample buffer (Novex; Invitrogen) and subsequently subjected to SDS-polyacrylamide gel electrophoresis (PAGE) on a 4–15% Mini Protean TGX precast gel (Bio-Rad Laboratories, Hercules, CA, USA).

### Purification of dsRNA–vRNP complex from infected cells

A549 cells were infected with the FLAG–PB2 virus at an MOI of 0.1 and incubated in MEM/BSA at 37°C. At 24 hpi, cells were scraped from dishes in ice-cold PBS and subsequently pelleted through centrifugation at 780 × *g* for 10 min at 4°C. The resultant pellets were resuspended in lysis buffer [50 mM Tris-HCl (pH 8.0), 150 mM NaCl, 5 mM MgCl2, 10% glycerol, 0.05% NP-40, 2 mM dithiothreitol (DTT), 10 mM ribonucleoside-vanadyl complex (New England Biolabs, Beverley, MA, USA), and 1× complete EDTA-free protease inhibitor (Roche, Mannheim, Germany)]. A rotation was implemented for 15 min at 4°C, followed by centrifugation at 20,000 × *g* for 15 min at 4°C. The resultant supernatant was incubated with anti-FLAG M2 affinity gel (Merck) on a rotator for 1 h at 4°C. The gels underwent a single wash with lysis buffer and three consecutive washes with wash buffer [50 mM Tris-HCl (pH 8.0), 200 mM NaCl, 50 mM Na2HPO4, and 2 mM DTT], subsequently eluting in wash buffer with 500 ng/µL FLAG peptide (Merck) through rotation on a rotator for 30 min at 4°C. The FLAG-tagged vRNP was further purified by glycerol gradient ultracentrifugation as described prior. Subsequently, the cellular lysates, eluates, and purified fractions were subjected to electrophoresis on an SDS-polyacrylamide gel, followed by silver staining or immunoblotting using anti-NP mouse monoclonal antibody (1:10000 dilution, ab20343; Abcam, Cambridge, UK) and anti-PB1 goat polyclonal antibody (1:10000 dilution, sc-17601; Santa Cruz Biotechnology) as primary antibodies. The secondary antibodies used were horseradish peroxidase-conjugated anti-mouse (NA931; GE Healthcare) and anti-goat (ab6741; Abcam) antibodies.

### Expression and purification of recombinant NS1 protein

The NS1 protein fused with a His-tag was expressed in *E. coli* Rosseta (DE3) pLysS cells using the pET-14b plasmid. The *E. coli* transformants were cultured at 37°C in LB medium supplemented with 100 µg/mL of ampicillin. Upon reaching an optical density at 600 nm (OD600) of 0.8, 1 mM of isopropylthio-β-galactoside (IPTG) was introduced, and the cells were grown for an additional 4 h at 37°C. Following this, *E. coli* cells were harvested by centrifugation at 8,000 × *g* for 5 min and resuspended in binding buffer [20 mM Tris-HCl (pH 8.0), 500 mM NaCl, and 5 mM imidazole]. The cells were lysed by sonication in an ice-cold water bath. Recombinant NS1 was purified using a Ni^2+^-immobilized column (His-Bind Purification Kit; Merck) according to the manufacturer’s instructions.

### *In vitro* RNA synthesis using virion-derived vRNPs

A concentration of 0.01 mg/mL of purified vRNP was subjected to incubation in 50 mM Tris-HCl buffer (pH 8.0) containing 5 mM MgCl2; 40 mM KCl; 1 mM DTT; 10 μg/mL actinomycin D; 1 mM each of ATP, CTP, GTP, and UTP; 1 U/μL RNasin Plus RNase inhibitor; and 1 mM ApG (IBA Lifesciences, Göttingen, Germany) at 30°C for 15 min. For the purpose of masking the dsRNA with NS1, 0.1 mg/mL of the purified recombinant NS1 was introduced into the *in vitro* transcribed vRNP mixture, followed by a 1-h incubation period on ice.

### HS-AFM

A 2 µL aliquot of the specimen was deposited onto freshly cleaved mica devoid of surface modification. Following incubation for 5 min at ambient temperature, the mica surface was thoroughly washed with imaging buffer [50 mM Tris-HCl (pH 8.0), 5 mM MgCl2, 40 mM KCl, and 1 mM DTT]. The mica surface was submerged in a liquid chamber filled with imaging buffer for observation using an HS-AFM system (Nano Explorer; Research Institute of Biomolecule Metrology Co., Ltd., Ibaraki, Japan). HS-AFM was executed at room temperature in the tapping mode, wherein the cantilever oscillated vertically at its resonant frequency during lateral and vertical scanning of the cantilever chip, intermittently tapping the sample surface. Images were captured at a rate of two images per second using cantilevers featuring a 0.1 N/m spring constant and a resonance frequency in water of 0.6 MHz (BL-AC10DS; Olympus, Tokyo, Japan). To acquire high-resolution images, electron-beam-deposited tips [49] were used. The *in situ* observation of RNA digestion by RNase was executed through introducing a 0.2 U/µL of ShortCut RNase III (New England Biolabs) to the AFM liquid cell during imaging. To facilitate the detachment of the NS1 protein from the dsRNA, the specimen underwent prolonged observation, gradually diminishing the setpoint on the AFM system. A minimum of five independent experiments were conducted, and all HS-AFM images were viewed and analyzed using Kodec 4.4.7.39 [50]. Individual images were subjected to a low-pass filter and flattening to eliminate spike noise and flatten the xy-plane, respectively.

### Statistics and reproducibility

All statistical analyses were conducted using the Prism 9.5.0 software (GraphPad, San Diego, CA, USA), with the specific methodology for each analysis delineated in the respective figure legends. A *p* value of less than 0.05 was considered statistically significant. Each experiment documented in the manuscript was replicated at least three times, yielding consistent and reproducible results.

## Data availability

All data substantiating the conclusions of this investigation are contained within the Manuscript and Supplementary Information files or can be obtained from the corresponding authors upon judicious and reasonable request.

## Acknowledgements

We thank Yoshihiro Kawaoka for providing us with plasmids, Takashi Fujita for giving us RIG-I KO 293T cells, and Noriyuki Kodera for fabricating electron-beam-deposited HS-AFM tips. We also thank Junichi Kajikawa for helpful discussion. We thank Editage (www.editage.com) for English language editing.

## Supporting information

**S1 Fig. dsRNA production in mock-infected cells.**

Cells subjected to mock infection were fixed at designated time points. After fixation, cells were stained with anti-NP and anti-dsRNA antibodies. Hoechst staining was used to stain cellular nuclei. The scale bars denote 20 µm.

**S2 Fig. Effect of MOI on IAV dsRNA production.**

Vero (A) and A549 (B) cells were infected with the IAV WSN strain at MOIs of 0.1 or 5 and subsequently fixed at 10 and 14 hpi, respectively. The presence of NP and dsRNA in infected cells was detected by IFA using anti-NP and anti-dsRNA antibodies, respectively. All dsRNA-producing cells within a given viewpoint are indicated by arrows. Hoechst staining was used to stain cellular nuclei. The scale bars represent 50 µm.

**S3 Fig. Simultaneous detection of NS1 and NEP in dsRNA-producing cells.**

A549 cells were infected with the IAV WSN strain at an MOI of 0.1 and subsequently fixed at 14 hpi. NS1 and NEP expression in dsRNA-producing cells were concomitantly detected by IFA using anti-NS1, anti-NEP, and anti-dsRNA antibodies. To confirm NP expression in dsRNA-producing cells, NS1 and NP were simultaneously detected using anti-NP antibodies. Cell nuclei were stained with Hoechst. Arrows indicate infected cells producing dsRNAs. The scale bars denote 20 µm.

**S4 Fig. Isolation of the vRNP–looped RNA complex from cells infected with FLAG– PB2 virus.**

vRNPs purified from FLAG–PB2 virus-infected A549 cells were subjected to SDS-PAGE followed by silver staining (A) and western blotting (B). The supernatant from cell lysis (Input), immunoprecipitate eluted with the FLAG peptide (IP), and purified fractions following ultracentrifugation (Nos.1–11) were examined. In panel A, bands corresponding to RNA polymerases (Pols) and NP are indicated by arrows. The verification of vRNP purification was conducted in panel B using anti-PB1 and anti-NP antibodies. Molecular weight marker proteins (M) were used as reference proteins. Uncropped gel and blot images are shown in S10 Fig. (C) Digestion of looped RNA by RNase III. Purified vRNP associated with looped RNA was observed using HS-AFM. During the observation, RNase III was introduced, and digestion of the looped RNA was monitored. Five representative images captured at designated intervals are presented, displaying digestion occurring between 396 and 397 s. Scale bars: 50 nm.

**S5 Fig. Generation of recombinant NS1 protein.**

(A) Expression of recombinant NS1 protein. *E. coli* cellular precipitates, both pre-induction (−) and post-induction (+) with IPTG, underwent SDS-PAGE followed by Coomassie Brilliant Blue (CBB) staining. The band corresponding to expressed NS1 is denoted by an arrow. (B) Purification of recombinant NS1 protein. *E. coli* expressing NS1 was subjected to sonication and centrifugation. The resulting precipitate (P) and supernatant (S) were analyzed via SDS-PAGE followed by CBB staining to ascertain the acquisition of recombinant NS1 as a soluble protein. Subsequently, the supernatant was subjected to purification using a Ni^2+^-immobilized column. After washing the column with Binding and Wash buffer, the recombinant NS1 was eluted using Elute buffer, as indicated by the arrow. Molecular weight marker proteins (M) were utilized for reference. The flow-through of the affinity column is labelled as FT. Uncropped gel images are shown in S10 Fig.

**S6 Fig. Cross-sectional analysis of the looped structure associated with vRNP.**

(A) Determination of the height of the vRNP and looped structures. A cross-sectional analysis was performed along the delineated red lines in the HS-AFM images (upper panels). The quantified heights of both vRNPs and looped structures are shown in the lower panels. (B) Quantification of the width of the looped structure. HS-AFM images analogous to that in Fig 3E (+NS1) are presented, and a cross-sectional analysis was conducted at distinct positions along the looped structure, as denoted by red lines. The measured widths are presented in the lower panels. All scale bars denote 50 nm.

**S7 Fig. Association between the localization of dsRNA and viral protein expression in ΔNS1 virus-infected cells.**

A549 cells were infected with ΔNS1 virus at an MOI of 0.1 and subsequently fixed at 24 hpi. dsRNA was detected by IFA using anti-dsRNA antibodies. Viral protein expression was examined using anti-NP (A), anti-NA (B), anti-M1 (C), and anti-NEP (D) antibodies. Cells exhibiting dsRNA production in the nucleus and cytoplasm are shown in the left and right panels, respectively. Arrows indicate cells producing dsRNA and concurrently expressing the designated viral proteins. Scale bars represent 10 µm.

**S8 Fig. dsRNA production and IRF3 nuclear translocation in wild-type IAV-infected cells.**

A549 cells were infected with the wild-type IAV WSN strain at an MOI of 0.2 and were subsequently fixed at 24 hpi. IFA was conducted to detect dsRNAs using anti-dsRNA antibodies. To examine the nuclear translocation of IRF3, anti-IRF3 antibodies were applied. Nuclear translocation of IRF3 was observed in certain dsRNA-producing cells (upper panels), whereas the other cells did not exhibit nuclear translocation of IRF3 (lower panels). Cell nuclei were stained with Hoechst. The scale bars represent 20 µm.

**S9 Fig. Nuclear translocation of IRF3 in RIG-I KO 293T cells transfected with poly(I:C).**

Wild-type 293T (upper panels) and RIG-I KO 293T (middle and lower panels) cells were transfected with poly(I:C). To restore RIG-I functionality, RIG-I KO 293T cells were transfected with RIG-I expression plasmid (pRIG-I) at 2 h post-poly(I:C) transfection (lower panels). Cellular fixation was performed at 24 hpt, followed by staining with anti-dsRNA and anti-IRF3 antibodies. Cell nuclei were stained with Hoechst. The scale bars represent 20 µm.

**S10 Fig. Uncropped gel and blot images.**

(A) Uncropped gel image for S4A Fig. (B) Uncropped blot image for S4B Fig (upper panel). (C) Uncropped blot image for S4B Fig (lower panel). (D) Uncropped gel image for S5A Fig. (E) Uncropped gel image for S5B Fig.

**S1 Table. Number of dsRNA-producing cells infected with wild-type IAV WSN strain at different MOI.**

**S2 Table. Number of viral protein expressing and dsRNA-producing A549 cells infected with wild-type IAV WSN strain at MOI of 0.1.**

**S1 Movie. Digestion of the looped RNA by RNase III.**

RNase III was added during HS-AFM observation of looped RNA–vRNP complex. Scan area: 300×300 nm^2^. Observation period: 90 sec.

**S2 Movie. Detachment of NS1 from the looped RNA.**

The looped structure observed after NS1 addition was observed over an extended duration using HS-AFM with applying an augmented force. Scan area: 300×300 nm^2^. Observation period: 450 sec.

